# A transition from SoxB1 to SoxE transcription factors is essential for progression from pluripotent blastula cells to neural crest cells

**DOI:** 10.1101/359752

**Authors:** Elsy Buitrago-Delgado, Elizabeth N. Schock, Kara Nordin, Carole LaBonne

## Abstract

The neural crest is a stem cell population unique to vertebrate embryos that gives rise to derivatives from multiple embryonic germ layers. The molecular underpinnings of potency that govern neural crest potential are highly conserved with that of pluripotent blastula stem cells, suggesting that neural crest cells may have evolved through retention of aspects of the pluripotency gene regulatory network (GRN). A striking difference in the regulatory factors utilized in pluripotent blastula cells and neural crest cells is the deployment of different subfamilies of Sox transcription factors; SoxB1 factors play central roles in the pluripotency of naïve blastula and ES cells, whereas neural crest cells require SoxE function. Here we explore the shared and distinct activities of these factors to shed light on the role that this molecular hand-off of Sox factor activity plays in the genesis of neural crest and the lineages derived from it. Our findings provide evidence that SoxB1 and SoxE factors have both overlapping and distinct activities in regulating pluripotency and lineage restriction in the embryo. We hypothesize that SoxE factors may transiently replace SoxB1 factors to control pluripotency in neural crest cells, and then poise these cells to contribute to glial, chondrogenic and melanocyte lineages at stages when SoxB1 factors promote neuronal progenitor formation.

## Introduction

The neural crest is a major evolutionary innovation of vertebrates, and acquisition of these cells allowed for the generation of many vertebrate-specific features (Hall, 2009; Hall., 2013; Le Douarin et al., 2008). First described by Wilhelm His 150 years ago, neural crest cells are distinguished by their ability to contribute to the vertebrate body plan a diverse array of cell types associated with multiple germ layers (His, 1868; Le Douarin et al., 2008). Referred to as the “Zwischenstrang” by His (His, 1868). The evolutionary acquisition of these cells allowed for the formation of a myriad of novel structures, including a “new head”, to be layered onto the simple chordate body plan (Hall, 2009). Within developing embryos, neural crest cells retain broad multi-germ layer potential even as neighboring cells become lineage restricted (Prasad et al 2012). Ultimately, neural crest cells give rise to a diverse array of derivatives that includes chondrocytes, melanocytes, and neurons and glia of the peripheral nervous system (Le Douarin et al., 2008; Taylor and LaBonne, 2007). While much has been learned about the signaling pathways and transcriptional responses required for formation of the neural crest and the subsequent lineage diversification of these cells, how specific factors contribute to the broad developmental potential of neural crest cells remains unclear.

A core network of transcriptions factors controls pluripotency in the blastula, including members of the Sox (SRY-related high mobility group (HMG)-box) family of transcription factors (Takahashi and Yamanaka, 2006). Sox proteins are highly conserved and contain a DNA-binding domain known as the HMG-box. Based upon homology within the HMG domain and other structural regions of the protein, Sox factors are divided into nine subfamilies (Bowles et al., 2000; Julian et al., 2017; Schepers et al., 2002). Two subfamilies of particular importance for pluripotency in blastula and neural crest cells are SoxB1 and SoxE factors, respectively. SoxB1 factors (*Sox1/2/3*) are essential regulators of the stem cell state in both blastula cells and ES cells (Guth and Wegner, 2008; Sarkar and Hochedlinger, 2013; Kopp et al., 2008; Boer et al., 2007). SoxB1 proteins are maternally expressed and highly enriched in the pluripotent cells (inner cell mass) of the blastula, consistent with a role during the early stages of development (Avilion et al., 2003; Buitrago-Delgado et al., 2015). Additionally, SoxB1 proteins can act as transcriptional activators of genes essential for pluripotency. For example, SOX2 functions with OCT4, another core pluripotency factor, to initiate a gene regulatory network involved in maintaining a stem cell state (Abdelalim et., al 2014; Masui et al., 2007; Takahashi and Yamanaka, 2006). Subsequent to their functions in pluripotent blastula cells, expression of *SoxB1* factors becomes restricted to the prospective neuroectoderm, where they are essential for the establishment of a neural progenitor state (Graham et al., 2003; Guth and Wegner, 2008; Rex et al., 1997; Streit et al., 1997).

Neural crest cells and pluripotent blastula cells display remarkable similarity in their gene expression profiles (Buitrago et al., 2015), and in the signals and epigenetic mechanisms that control this gene expression (Geary and LaBonne, 2018; Rao and LaBonne, 2018). Many neural crest potency factors are first expressed in naïve blastula cells, including *Snail1, Myc, Foxd3, Ets1, Ap2*, and *Vent2* (Buitrago et al., 2015; Ochoa et al., 2012; Bellmeyer et al., 2003). A striking difference in the gene regulatory networks (GRNs) controlling pluripotency in blastula cells versus neural crest cells are the Sox factors deployed. In neural crest stem cells the essential Sox factors are the SoxE factors, rather than SoxB1 factors. SoxE proteins (Sox 8, 9, 10) play important roles in establishing the neural crest stem cell population, and subsequently direct the formation of a subset of neural crest cell lineages (Cheung and Briscoe, 2003; Haldin and LaBonne, 2010; Hong and Saint-Jeannet, 2005; Kim et al., 2003; Lefebvre et al., 2017; Spokony et al., 2002). For example, Sox9 is required for formation of chondrocytes, while Sox10 is essential for melanocyte and glial cell formation (Akiyama et al., 2002; Aoki et al., 2003; Bi et al., 1999; Chen et al., 2001; Haldin and LaBonne, 2010; Lee et al., 2011, Mori-Akiyama et al., 2003; Suzuki et al., 2006). While distinct roles for SoxE factors in directing these neural crest cell lineage decisions have been defined, little is understood about their contributions to maintaining the neural crest stem cell state. One hypothesis is that SoxE factors play an analogous role in the neural crest to that of SoxB1 factors in naïve blastula cells with to maintain a pluripotent state. However, this raises the question of why a hand-off from SoxB1 to SoxE factors is necessary and/or advantageous.

Here, we examine the activities of SoxB1 and SoxE factors in the neural crest and pluripotent blastula cells of early *Xenopus* embryos. We demonstrate that SoxB1 and SoxE factors exhibit both unique and redundant activities in these cell types. Our findings suggest a model in which the essential role of SoxB1 in regulating the blastula stem cell state is handed off to SoxE factors which function in part to retain the developmental potential of the neural crest.

## Results

### SoxB1 and SoxE factors are expressed sequentially during *Xenopus* development

We recently demonstrated that much of the regulatory network that controls the pluripotency of blastula stem cells/animal pole cells is shared with neural crest cells, shedding new light on the origins of neural crest cells and the evolution of vertebrates (Buitrago-Delgado et al., 2015). We found that factors that have long been considered neural crest potency factors are first expressed in blastula animal pole cells, and are required for the pluripotency of these cells; however, a subset of neural crest factors does not show prior expression in pluripotent blastula cells. These factors represent true evolutionary novelties whose co-option into the neural crest GRN may have played a key role in endowing neural crest cells with the ability to transform the vertebrate body plan. Prominent among these factors are the SoxE family transcription factors, Sox9 and Sox10.

We compared the expression of *SoxB1* and *SoxE* transcription factors from blastula to late neurula stages using *in situ* hybridization to better understand the transition in their expression. The SoxB1 factors *Sox2* and *Sox3* are robustly expressed in the animal pole region of blastula embryos, where pluripotent cells reside (Figure 1A). By contrast, expression of the SoxE factors *Sox9* and *Sox10* cannot be detected until late gastrula or early neurula stages respectively, when they mark the neural crest populations at the neural plate border (Figure 1B). A third SoxE factor, *Sox8*, is expressed at low levels in blastula stage embryos but is not detectable by the onset of gastrulation. By midgastrula stages (stage 11) *Sox8* expression can be detected in neural crest regions as previously reported (Hong and Saint-Jeannett 2005; O’Donnell et al., 2006) as well as in anterior prospective ectoderm regions where *Sox2* expression has been diminished (Figure 1B). By late gastrula/early neurula stages, expression of *Sox2* and *Sox3* has been restricted to the prospective neural plate, marking the transition from their role in pluripotency to their subsequent roles in maintaining neuronal progenitor cells. The expression of SoxB1 and SoxE factors overlap at late gastrula stages, when both are expressed in neural crest regions of the neural plate border; however, by early neurula stages their expression is distinct. At this time *SoxE* factors mark the neural crest (and in the case of *Sox9* and *Sox10* also the otic placode), and *SoxB1* factors mark the neural plate and preplacodal region (Figure 1A, B).

**Figure 1.**
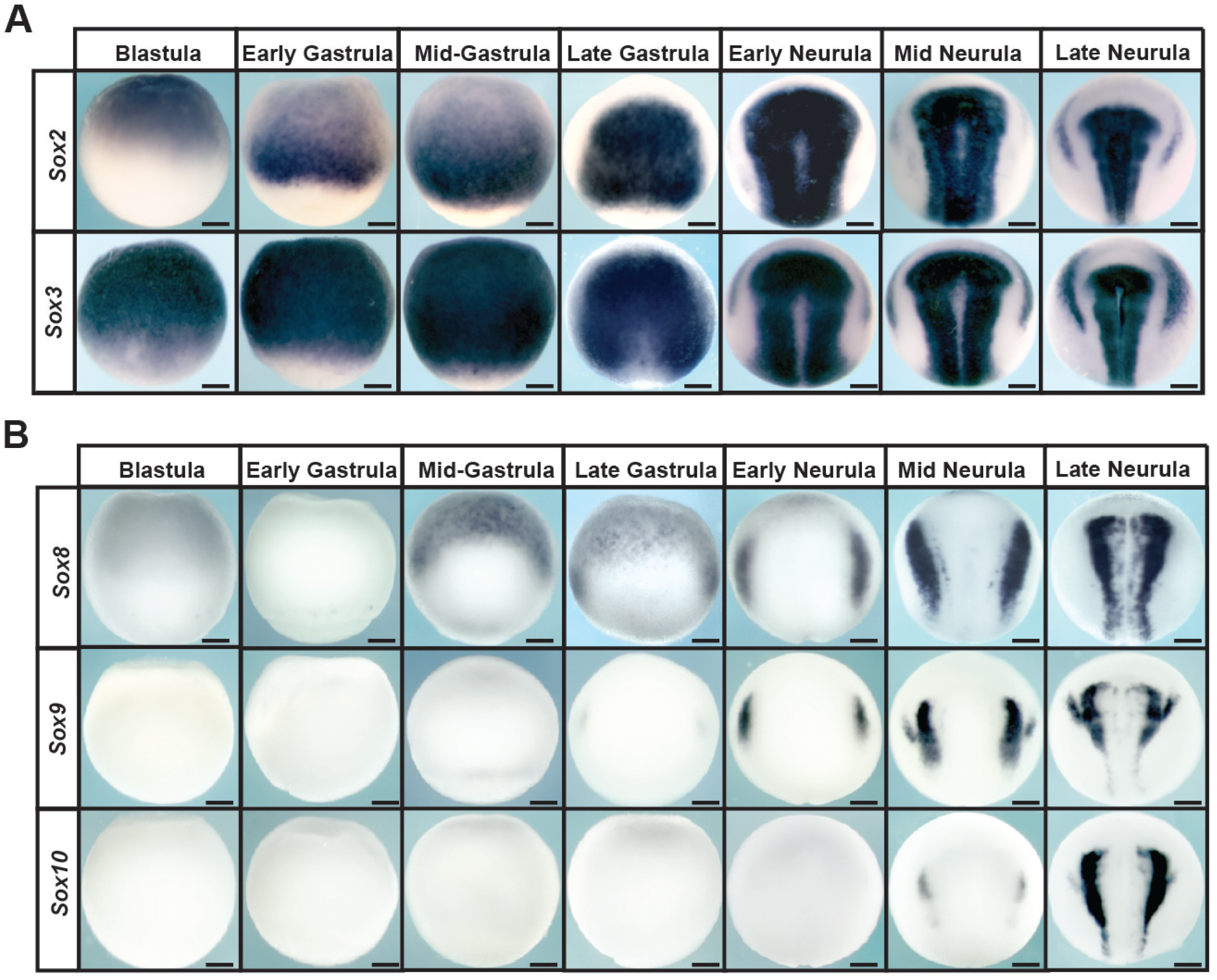
Expression of *SoxB1* and *SoxE* factors in *Xenopus* embryos. (A) *In situ* hybridization examining *Sox2* and *Sox3* expression in wildtype *Xenopus* embryos collected between blastula and late neurula stages. (B) *In situ* hybridization examining *Sox8, Sox9*, and *Sox10* expression in wildtype *Xenopus* embryos collected between blastula and late neurula stages.

### Premature SoxE activity interferes with proper pluripotency gene expression

The distinct deployment of SoxB1 and SoxE factors in pluripotent blastula cells and neural crest respectively suggests that these proteins may have distinct activities that favor their function in these different contexts. In particular we were interested in whether exclusion of SoxE factors from pluripotent blastula cells was essential to maintain a stem cell state. To determine this, mRNA encoding Sox9 or Sox10 was injected in once cell of two cell embryos. Injected embryos were cultured to blastula stages when expression of four key genes expressed in pluripotent cells, *Oct25, Vent2, Id3* and *TF-AP2*, was examined by *in situ* hybridization. Premature activity of either Sox9 or Sox10 led to strong down-regulation of these genes on the injected side of the embryo (Figure 2A, S1A). This result suggested that SoxE function at these stages may be incompatible with normal development of pluripotent blastula cells. In parallel experiments, Sox2 or Sox3 were expressed in blastula cells at protein levels matched to Sox9 and Sox10. While this too led to disruptions in gene expression, these changes were less pronounced than those observed in response to equivalent levels of Sox9 or Sox10 (Figure 2A, S1A). The changes in blastula gene expression in response to up-regulation of SoxB1 factors is consistent with findings in mouse and human ES cells, where both increases and decreases in Sox2 activity can lead to loss of pluripotency (Kopp et al., 2008; Boer et al., 2007; Yamaguchi et al., 2011; Takahashi &Yamanaka, 2006; Thomson et al, 2011). These findings suggest that pluripotent blastula cells, while sensitive to changes in the levels of Sox protein activity, are more sensitive to increases in the activity of SoxE family transcription factors.

**Figure 2.**
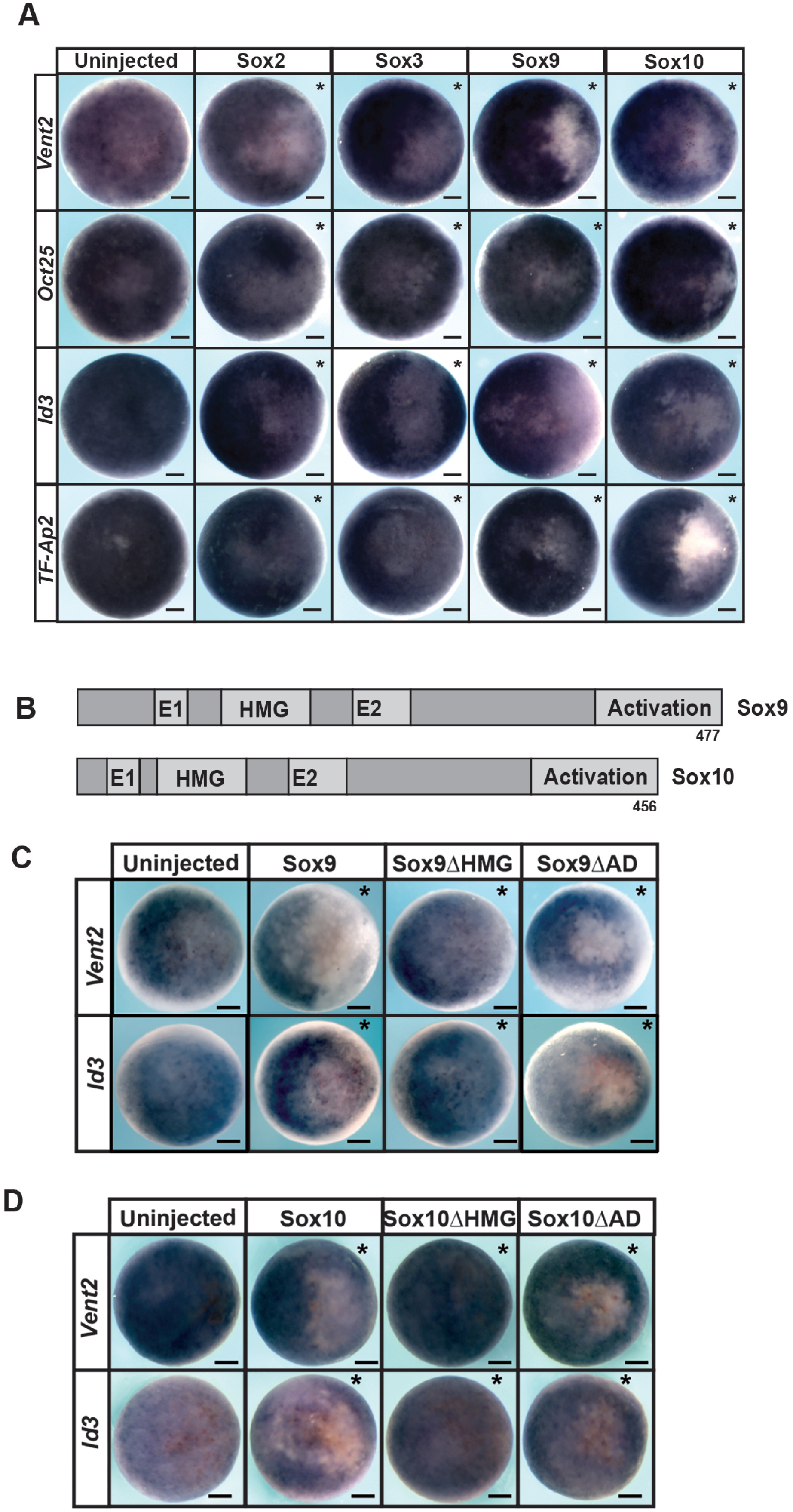
SoxE factors interferance with pluripotency gene expression depends upon DNA-binding capabilities. (A) *In situ* hybridization examining *Vent2, Oct25, Id3*, and *TF-AP2* in blastula stage (stage 9) embryos injected unilaterally with Sox2, Sox3, Sox9 or Sox10. (B) Schematic of Sox9 and Sox9 protein domains. (C) *In situ* hybridization examining *Vent2* and *Id3* expression in blastula stage (stage 9) embryos injected unilaterally with wildtype or mutant forms of Sox9. (D) *In situ* hybridization examining *Vent2* and *Id3* expression in blastula stage (stage 9) embryos injected unilaterally with wildtype or mutant forms of Sox10. Asterisk (*) denotes injected side of the embryo as marked by staining of the lineage tracer β-galactosidase (red).

We next asked if the down-regulation of blastula pluripotency genes in response to Sox9 or Sox10 was mediated by a specific domain of these SoxE proteins. We use mutant forms of these proteins in which either the activation domain or the HMG DNA binding domain had been deleted (Figure 2B). These mutant proteins were expressed at levels equivalent to the wild type protein as determined by western blot, and their ability to down-regulate *Id3* or *Vent2* expression at blastula stages was compared. We found that mutants lacking the activation could still down-regulate *Vent2* and *Id3* (Figure 2C,D). By contrast, Sox9 and Sox10 mutants lacking the HMG domain had no effect on gene expression in pluripotent cells. (Figure 2 C,D, S1B), suggesting that the observed down-regulation of pluripotency genes by SoxE factors is dependent upon DNA binding.

### SoxB1 and SoxE factors have distinct effects on neural crest and epidermis at neurula stages

Given that premature expression of SoxE factors in pluripotent blastula cells disrupts normal gene expression, we wished to determine if prolonged expression of SoxB1 factors in the cells that will become neural crest has effects distinct from SoxE activity. Importantly, however, comparing the effects of SoxB1 and SoxE activity in neural crest cells necessitates bypassing their effects on blastula cells. To accomplish this we expressed glucocorticoid receptor fusion proteins of both SoxB1 (*Sox2* and *Sox3*) and SoxE factors (*Sox9* and *Sox10*). These fusion proteins remain functionally inactive until embryos are treated with dexamethasone, allowing temporal control of their function in the developing embryo. Embryos were injected with mRNA encoding these factors in one cell at the two-cell stage. Injected embryos were treated with dexamethasone at stage 10 and cultured to neurula stages when they were harvested for analysis by *in situ* hybridization (Figure 3A). Interestingly, we found that inducing SoxB1 activity at the neural plate border at these stages led to down-regulation of neural crest factors *Foxd3* and *Snail2* (Figure 3B, S1C). By contrast, when SoxE proteins were similarly activated they enhance the expression of these neural crest markers (Figure 3B, S1C) consistent with previous results (Aoki et al.,2003; Saint-Germain et al., 2004; Taylor and LaBonne, 2005). These findings demonstrate that SoxE and SoxB1 activity have very different effects on the expression of neural crest markers at the neural plate border that are consistent with the spatiotemporal expression of these factors. To determine if these functional differences correlate with developmental timing (gastrula/neurula stage ectoderm) or cell type (neural crest) we also compared the effects of SoxB1 and SoxE activation on expression of the epidermal marker *Epidermal keratin*. Activation of either SoxB1 or SoxE factors downregulated the expression of this epidermal marker (Figure 3 C, S1C) suggesting that in this context, SoxB1 and SoxE factors have similar activities.

**Figure 3.**
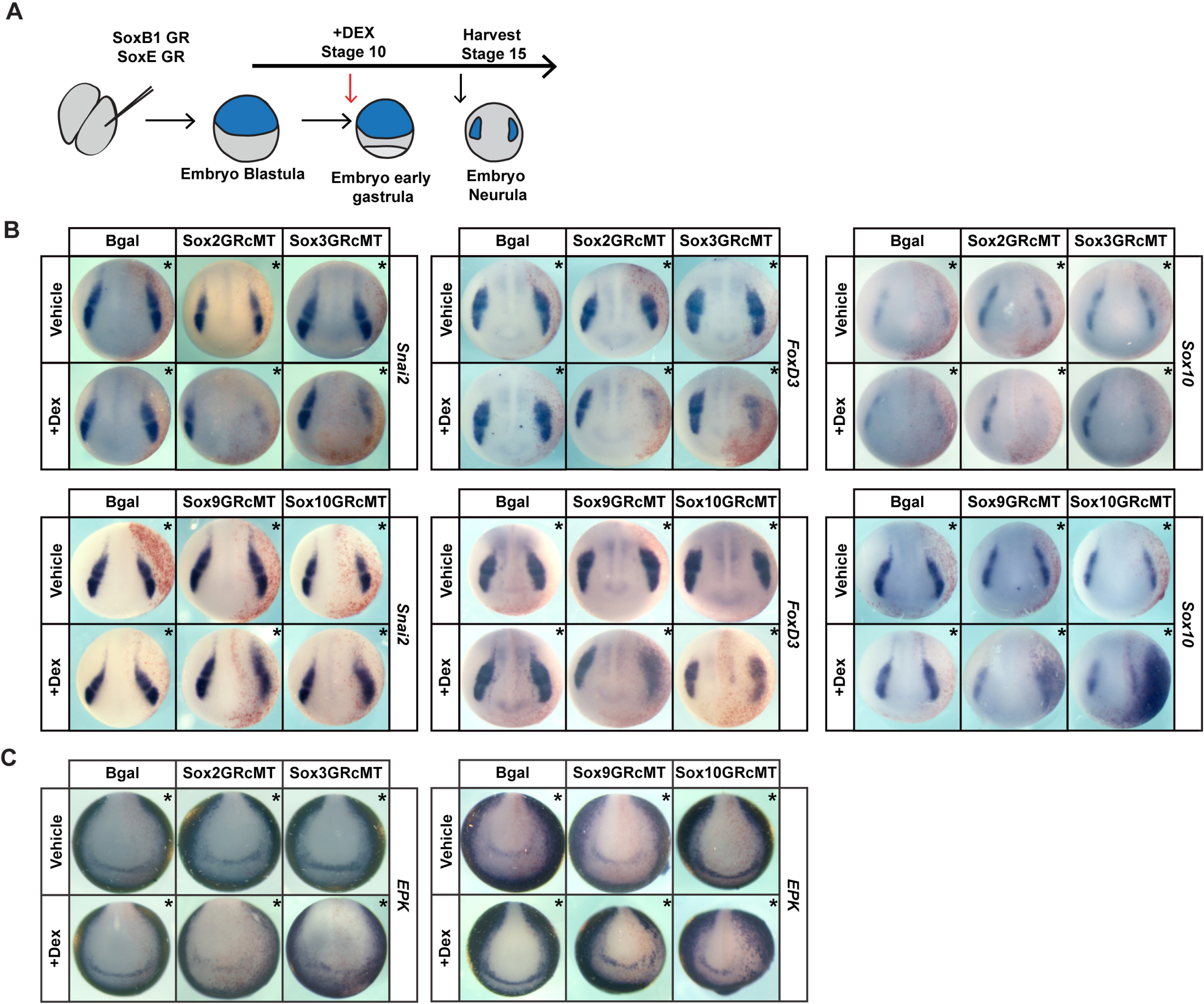
Inducing SoxB1 and SoxE protein activities at gastrulation leads to differential effects on neural crest formation, but similar effects on the ectoderm. (A) Schematic of experimental overview. (B) *In situ* hybridization assaying expression of neural crest factors *Snai2, FoxD3*, and *Sox10* in neurula (stage 15) embryos injected with inducible Sox2, Sox3, Sox9, or Sox10 constructs. (C) *In situ* hybridization assaying the expression of epidermal factor *EPK* in neurula (stage 15) embryos injected with inducible Sox2, Sox3, Sox9, or Sox10 constructs. Asterisk (*) denotes injected side of the embryo as marked by staining of the lineage tracer β-galactosidase (red).

### Increased SoxB1 or SoxE activity interferes with pluripotency

Given the observed down-regulation of key potency genes in pluripotent blastula cells, we examined the consequences of SoxE and SoxB1 activation on pluripotency. The animal pole cells of blastula stage embryos (state 8-9) are pluripotent, and explants of these cells can give rise to all cell derived from all three germ layers given proper instruction (Ariizumi & Asashima, 2001). For example, treatment of animal pole cells with activin instructs them to adopt a mesoderm or endoderm fate in a dose dependent manner (Asashima et al., 1990a; Asashima et al., 1990b; Lamb et al., 1993; Sasai et al., 1995). Using this system, we asked whether increased SoxE or SoxB1 activity would interfere with the ability of animal pole explants to adopt these fates in response to activin. Embryos were injected with mRNA encoding Sox2, Sox3 Sox9, or Sox10 in both cells at the 2-cell stage. Explants were manually dissected at blastula stage 8, and cultured with or without activin until early gastrula stages (stage 11.5) (Figure 4A). While control explants showed robust expression of the mesodermal marker *Brachyury* in response to low doses of activin, overexpression of SoxB1 or SoxE factors interfered with this response (Figure 4B, S1D) Similarly, SoxB1 or SoxE overexpression interfered with induction of endoderm in response to high doses of activin, as evidenced by loss of *Sox17* expression (Figure 4C, S1D). Thus, consistent with effects on pluripotency gene expression, up-regulation of Sox activity interfered with the functional pluripotency in these cells, and therefore in this context SoxB1 and SoxE factors exhibit similar activities.

**Figure 4.**
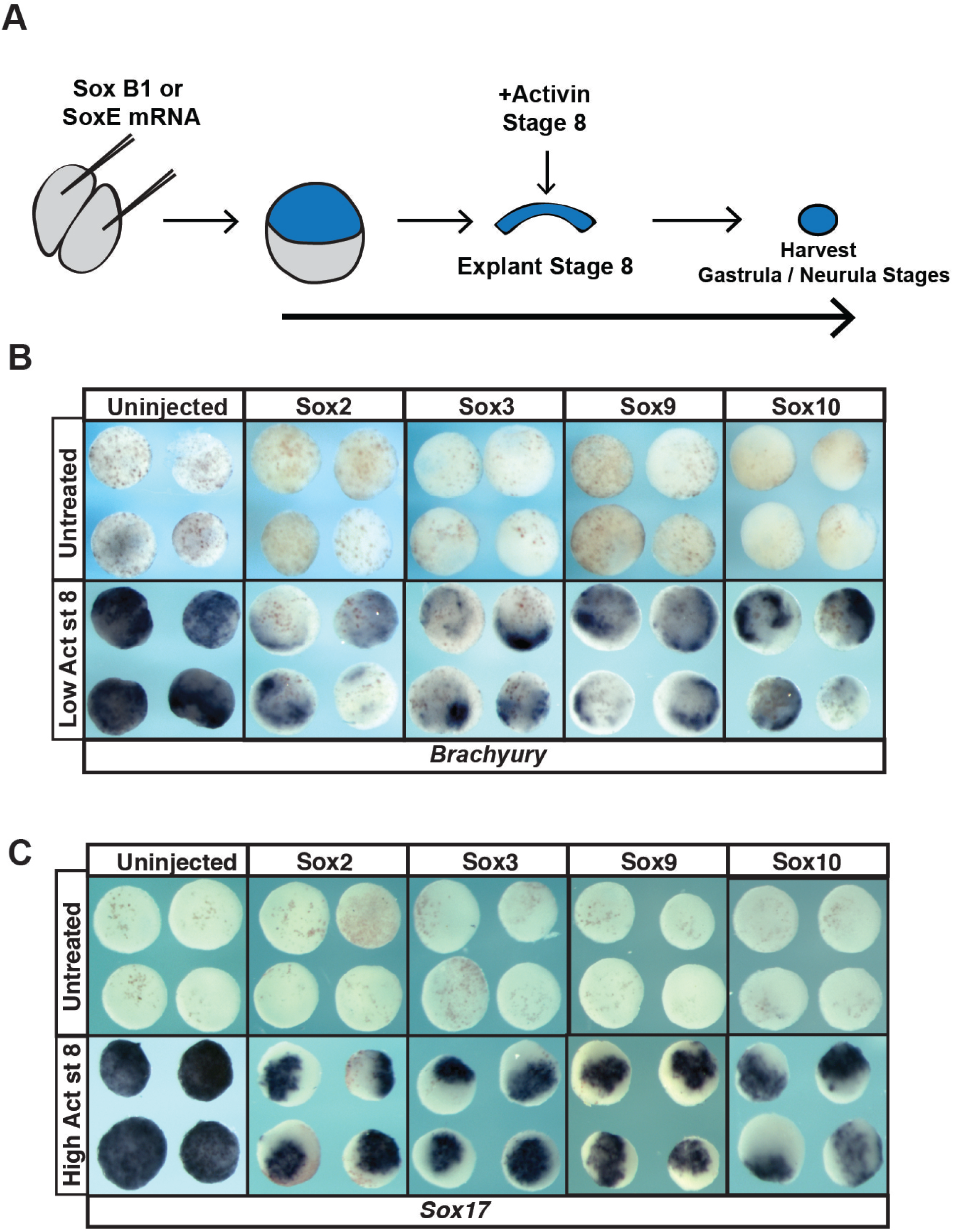
Overexpression of SoxB1 or SoxE factors impairs the formation of mesoderm and endoderm in animal pole explant assays. (A) Schematic of experimental overview. (B) *In situ* hybridization examining *Brachyury* expression in animal pole explants injected with Sox2, Sox3, Sox9 or Sox10 mRNA and cultured with or without a low dose of activin. (C) *In situ* hybridization examining *Sox17* expression in animal pole explants injected with Sox2, Sox3, Sox9 or Sox10 mRNA and cultured with or without a high dose of activin.

### Both SoxB1 and SoxE1 proteins can maintain pluripotency

The ability of the core pluripotency factors, including Sox2, to maintain developmental potential is concentration dependent (Kopp et al., 2008; Boer et al., 2007; Yamaguchi et al., 2011), and this may explain the diminished ability of animal pole explants to form mesoderm or endoderm when SoxB1 or SoxE factors are overexpressed. To more rigorously compare the ability of SoxB1 and SoxE factors to mediate pluripotency we carried out molecular replacement experiments. We used previously characterized translation blocking morpholinos targeting Sox2 and Sox3 to deplete the function of SoxB1 in animal pole cells (Schlosser et al., 2008). Cells depleted of SoxB1 factors are no longer competent to form mesoderm or endoderm in response to activin treatment, as assayed by expression of *Brachyury* and *Endodermin* (Figure 5A-C), confirming that SoxB1 function is essential for pluripotency in blastula stem cells. Interestingly, the ability of these cells to adopt mesodermal or endodermal states could be rescued not only by expressing Sox2 or Sox3 but also by expressing Sox9 or Sox10 (Figure 5B-C, S1). Thus, although they are not normally expressed in pluripotent cells of the blastula, Sox9 and Sox10 do have the ability to maintain pluripotency, although they may do so less robustly than SoxB1 factors do.

**Figure 5.**
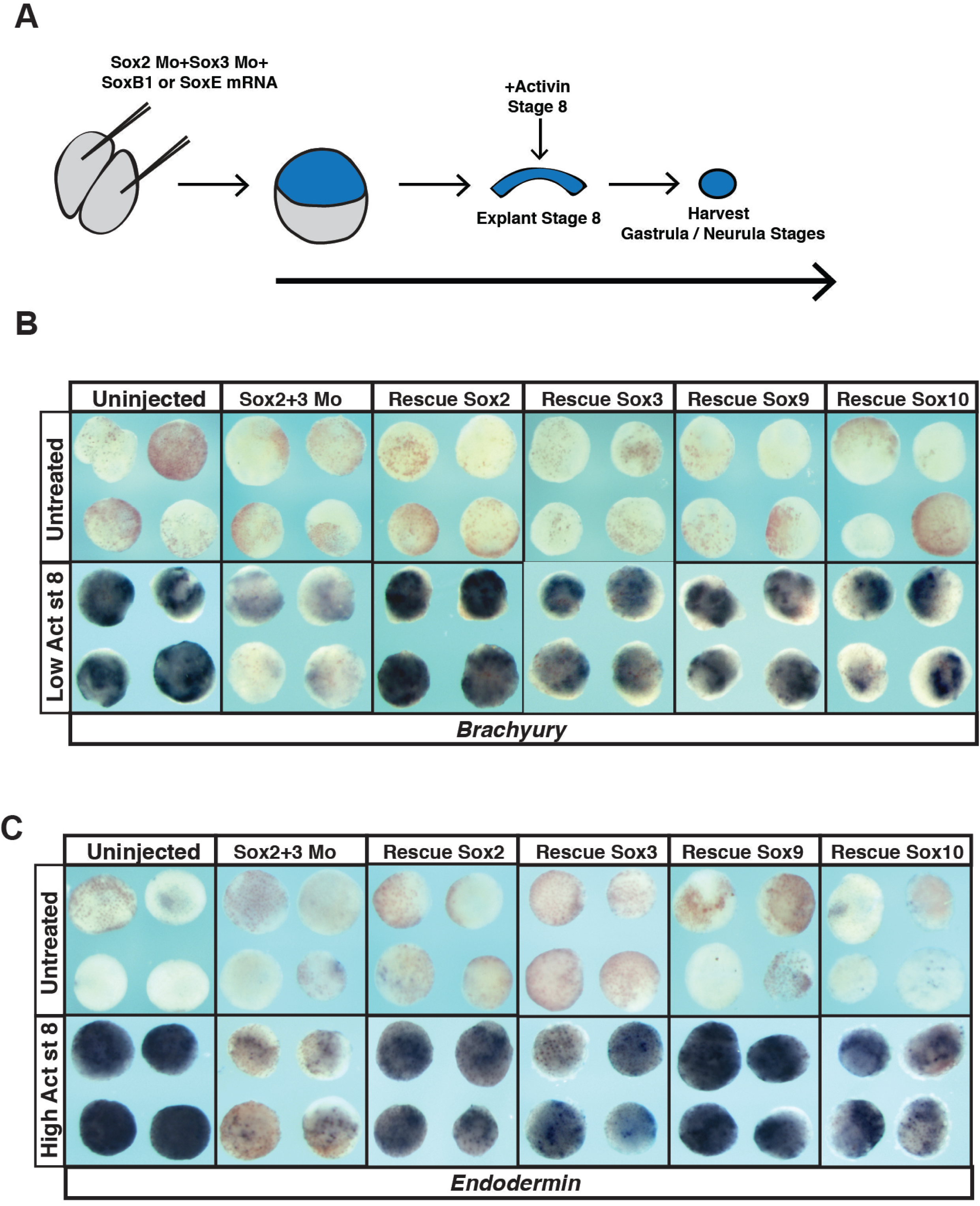
SoxB1 and SoxE factors can both restore potential to animal pole explants following morpholino-mediated depletion of Sox2 and Sox3. (A) Schematic of experimental overview. (B) *In situ* hybridization examining *Brachyury* expression in animal pole explants injected with Sox2 and Sox3 morpholino and either Sox2, Sox3, Sox9 or Sox10 mRNA that were cultured with or without a low dose of activin. (C) *In situ* hybridization examining *Endodermin* expression in animal pole explants injected with Sox2 and Sox3 morpholino and either Sox2, Sox3, Sox9 or Sox10 that were cultured with or without a high dose of activin.

### SoxB1 proteins, but not SoxE proteins, can maintain a neuronal progenitor state

The ability of SoxE factors to replace SoxB1 factors in maintaining pluripotency raises the question of why a hand-off from SoxB1 to SoxE factors as cells progress from blastula stem cells to neural crest cells is necessary or advantageous. Given their later expression patterns, we hypothesized that these two families of Sox transcription factors might differentially poise cells for distinct lineage decisions. For example, subsequent to their roles in maintaining pluripotency in blastula stem cells, SoxB1 factors become restricted to the neural plate and are essential for formation of neuronal progenitor cells. Thus we wondered if SoxE factors could replace the function of SoxB1 factors in promoting a neural progenitor state. Animal pole explants can be induced to adopt a neural state in response to the BMP antagonist chordin. We first showed that morpholino-mediated depletion of Sox2 and Sox3 prevented chordin-mediated neural induction (Figure 6A-B). We then compared the ability of SoxB1 and SoxE factors to rescue the adoption of a neural state. Explants from morphant embryos co-injected mRNA encoding Sox2 or Sox3 expressed the neural marker *Sox11* in response to Chordin (Figure 6D-F, S1E). By contrast, neither Sox9 nor Sox10 equivalently rescued neural induction (Figure 6E-F, S1E). This indicates that SoxB1 factors posses a great ability to promote neural progenitor formation than do SoxE factors, consistent with their deployment in these cells at neurula stages.

**Figure 6.**
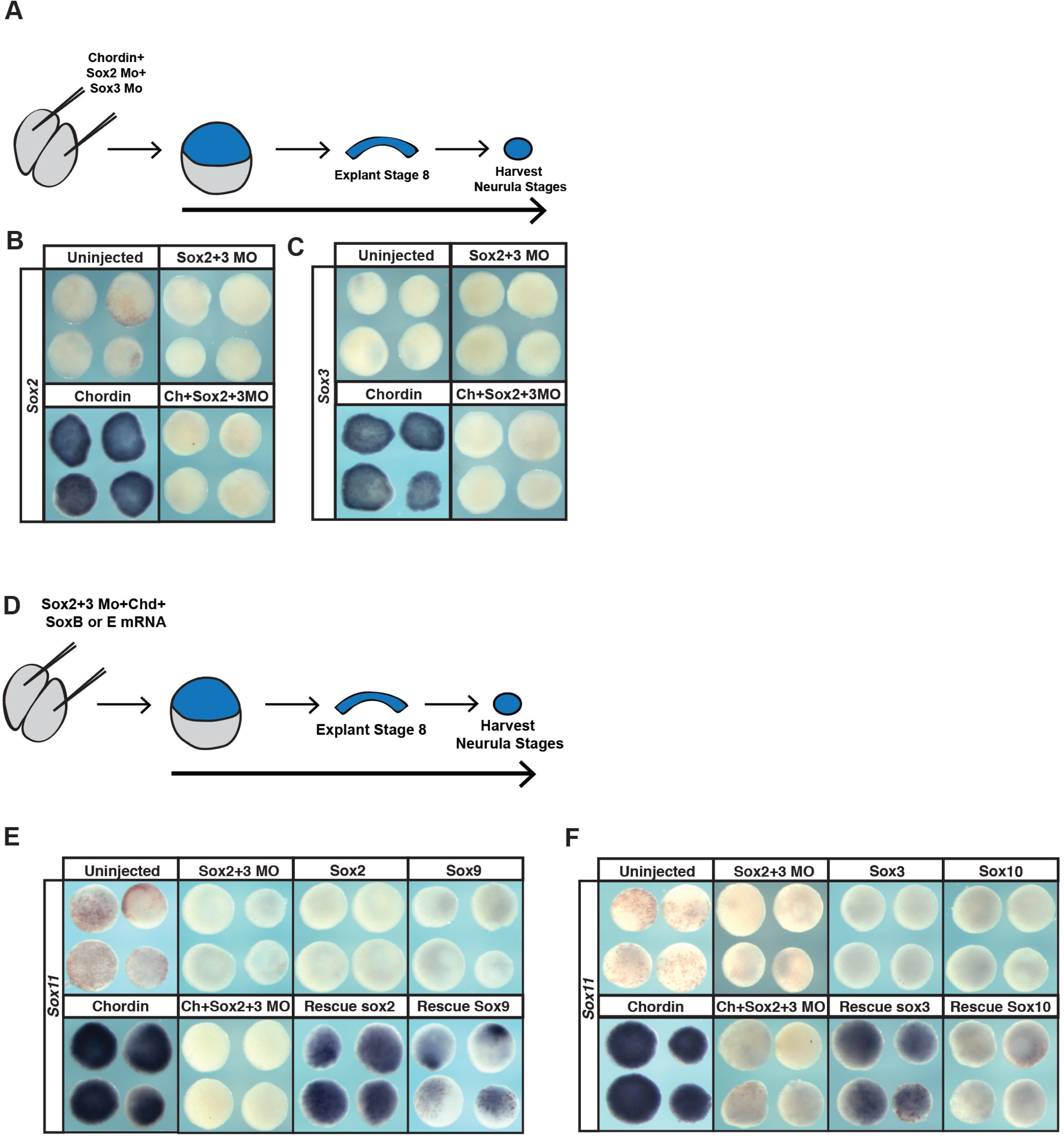
SoxE factors cannot fully replace the function of SoxB1 factors in animal pole explants induced to a neuronal lineage. (A) Schematic of experimental overview for knockdown experiments. (B) *In situ* hybridization examining *Sox2* expression in animal pole explants injected with Sox2 and Sox3 morpholino and chordin. (C) *In situ* hybridization examining Sox3 expression in animal pole explants injected with Sox2 and Sox3 morpholino and chordin. (D) Schematic of experimental overview for rescue experiments. (E) *In situ* hybridization examining *Sox11* expression in animal pole explants injected with Sox2 and Sox3 morpholino, chordin, and either Sox2 or Sox9 mRNA. (F) *In situ* hybridization examining *Sox11* expression in animal pole explants co-injected with Sox2 and Sox3 morpholino, chordin, and either Sox3 or Sox10.

### SoxE proteins, but not SoxB1 proteins, can promote the neural crest state

Given these findings, we next tested whether SoxB1 factors could replace SoxE functions in establishing a neural crest state. Animal pole cells can be reprogrammed to a neural crest state by expression of Wnt8 and Chordin (LaBonne & Bronner-Fraser, 1998, Figure 7A-E). Sox10 is required for establishing the neural crest state (Aoki et al., 2003; Honoré et al., 2003; Taylor & LaBonne, 2005) and morpholino-mediated depletion of Sox10 prevents expression of the neural crest markers *Sox9, Sox10* and *Foxd3* in these explants (Figure 7A, B, S2) Co-expression of either Sox9 or Sox10 could rescue the induction of the neural crest marker *FoxD3* (Figure 7D,E, S1E, S2) in these explants. By contrast Sox2 or Sox3 showed little or no ability to rescue *Foxd3* expression. These findings suggest that SoxB1 factors have only a limited ability to replace SoxE function in promoting formation and maintenance of neural crest cells.

**Figure 7.**
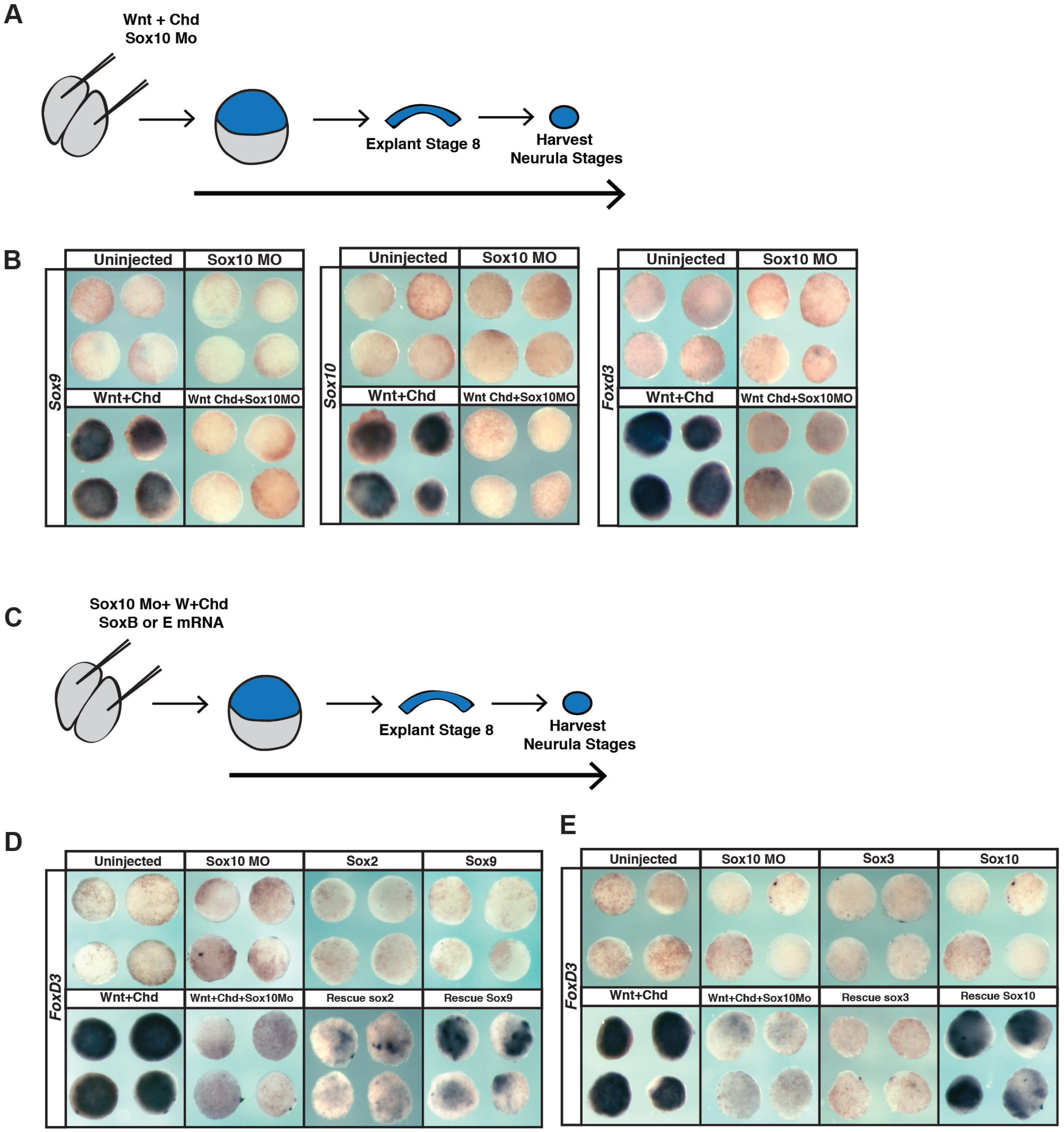
SoxB1 factors cannot replace the function of SoxE factors in animal pole explants induced to a neural crest state. (A) Schematic of experimental overview for knockdown experiments. (B) *In situ* hybridization examining *Sox9, Sox10*, or *FoxD3* expression in animal pole explants injected with Sox10 morpholino, Wnt8, and chordin. (C) Schematic of experimental overview for rescue experiments. (D) *In situ* hybridization examining *FoxD3* expression in animal pole explants injected with Sox10 morpholino, Wnt8, chordin, and either Sox2 or Sox9 mRNA. (E) *In situ* hybridization examining *FoxD3* expression in animal pole explants injected with Sox10 morpholino, Wnt8, chordin, and either Sox3 or Sox10.

## Discussion

One hundred and fifty years after the discovery of the neural crest, the primary synapomorphy of vertebrates, by Wilhelm His in 1868, much remains to be learned about these cells. The neural crest is distinguished by its retention of stem cell attributes long past the time when neighboring cells in the early embryo have undergone lineage restriction, as well as by the broad and diverse set of derivatives to which these cells ultimately contribute. While there is considerable overlap in the GRNs controlling pluripotency in blastula stem cells and neural crest cells (Buitrago et al., 2015), a major difference between these cell types is the sub-type of Sox family transcription factors deployed (Figure 1).

A dramatic change in the expression of SoxB1 and SoxE factors is observed as embryos progress from early cleavage and blastula stages, a time in development when populations of pluripotent cells are present, to neurula stages when significant lineage restriction has occurred, and definitive neural crest cells are present. The SoxB1 factors, *Sox2* and *Sox3*, are highly expressed in early pluripotent cells, but become restricted to the medial neural plate following gastrulation as animal pole cells become progressively lineage restricted (Fig. 1A). By contrast, expression of the SoxE factors *Sox9* and *Sox10* is absent from naïve blastula cells but becomes up-regulated at the neural plate border (NPB) by late gastrula stages, making the definitive neural crest stem cell population (Fig. 1B). By early neurula stages the expression domains of SoxB1 factors and SoxE factors have become mutually excusive. This is likely due, at least in part, to the repressive activity of Snail2 on *Sox2* expression (Acloque et al., 2011). The sequential deployment of different sub-families of Sox factors is reminiscent of what has been observed for Fox family transcription factors (Charney et al., 2017; Xu et al., 2009), suggesting that the SoxB1 to SoxE transition could serve as a paradigm for understanding the sequential utilization of related transcription factors during development. Furthermore, these findings suggest that with respect to the evolution of neural crest, the co-option of SoxE factors into the GRN represents one of the true novelties that correlated with or drove the acquisition of this cell type at the base of the vertebrates.

Why might it be important to deploy distinct sub-families of Sox transcription factors to maintain pluripotency in neural crest cells versus naïve blastula cells? One clue may be derived from the cells types that deploy SoxB1 and SoxE factors during later events in embryogenesis. By neural plate stages the expression of SoxB1 factors becomes restricted to neural progenitor cells where they play an essential role in maintaining the neural progenitor state (Bergsland et al., 2011). Neural crest cells retain their broad developmental potential through neurulation, and the onset of migration (Light et al., 2011), before ultimately differentiating and contributing to a broad set of derivative cell types (Taylor and LaBonne, 2007; Prasad et al., 2012). At these later stages, SoxE factors function to promote the formation of a subset of neural crest derivatives. For example, Sox9 plays a key role in the formation of cartilage, whereas Sox10 is essential for the formation of melanocytes and glia (Akiyama et al., 2002; Aoki et al., 2003; Britsch et al., 2001, Lee et al., 2011; Taylor and LaBonne, 2005). The restricted expression of SoxB1 versus SoxE factors suggests that as cells exit from pluripotency, SoxB1 function might be better suited to promote the establishment of a neural progenitor state, whereas SoxE function better poises cells to adopt non-neuronal neural crest states such as cartilage, glia and melanocytes. Consistent with important sub-functionalization, we find that SoxE factors promote the neural crest cell state whereas SoxB1 factors inhibit the formation of neural crest cells (Figure 3B, 7D).

Interestingly, during neural differentiation of embryonic stem cell cultures, SoxB1 factors take part in an additional relay event (Wegner, 2011). Following SoxB1-mediated maintenance of the neural progenitor state, SoxC family transcription factors (Sox4/11/12) function to promote neural differentiation (Bergsland et al., 2006; Bergsland et al., 2011; Hoser et al., 2008). It has been proposed that in this context Sox2 functions as a pioneer factor to both activate pluripotency genes and poises neural precursor genes. As cells assume a neuronal progenitor state, Sox3 replaces Sox2 occupancy of neural progenitor genes and poises neuronal differentiation genes for later expression. Finally, during neuronal differentiation, the SoxC protein, Sox11, replaces Sox2/3 and promotes expression of neural differentiation targets, including *Lhx2, Pax2*, and *Tubb3* (Bergsland et al., 2011). Our results show that the SoxB1 factors, Sox2 and Sox3, can rescue chordin-mediated neural induction to a much greater degree than can Sox9 and Sox10 (Fig. 6E, F). By contrast SoxE factors can rescue Wnt/chordin-mediated neural crest induction more potently than can SoxB1 factors (Fig. 7D, E). We hypothesize that SoxB1 and SoxE factors can occupy an overlapping set of regulatory elements on target genes, and may assemble distinct regulatory complexes in some contexts. Precedence for this is found in the regulation of neural crest derived oligodendrocytes, where both Sox10 and Sox2 can bind the regulatory elements for myelin binding protein, but with different transcriptional outputs (Hoffmann et al., 2014). Going forward it will be important to examine the dynamics of SoxB1 and SoxE protein occupancy across the genome as cells progress from a naïve blastula to a neuronal progenitor or neural crest state.

Sox factors are a highly versatile family of transcription factors that play multiple reiterative roles at different stages during development (Akiyama et al., 2002; Kim et al., 2003; Sarkar and Hochedlinger, 2013). Post-translational modification can further contribute to the functional versatility of these factors, for example SoxE proteins can functions as activators or repressors depending upon whether they have been modified by Sumoylation (Lee et al., 2012; Taylor and LaBonne 2005). SUMOylated SoxE factors recruit transcriptional repressors to inhibit genes important for melanogenesis, including *Dct* (Lee et al., 2012). SoxE function can also be tuned by context dependent interactions with the SoxD factor, Sox5, which enhances SoxE-mediated activation of cartilage genes such as *Col2a1*, but inhibits SoxE mediated activation of melanocyte and glial specific genes (Lefebvre et al., 1998; Nordin & LaBonne, 2014; Stolt et al., 2006; Stolt et al., 2008). Thus, multiple distinct mechanisms could contribute to tuning SoxB1 function for pluripotency and neuronal progenitor functions, and SoxE factors for neural crest progenitors and glial, melanocyte and cartilage fates.

Consistent with other studies, we found that levels of Sox proteins expressed were an important determinant of their function. In mouse and human ES cells, either increased or decreased Sox2 expression leads to loss of pluripotency (Kopp et al., 2008; Boer et al., 2007; Yamaguchi et al., 2011; Takahashi &Yamanaka, 2006; Thomson et al, 2011). Similarly we found that increased levels of Sox2 or Sox3 could inhibit gene expression in pluripotent blastula cells, although these cells appeared more sensitive to increases in SoxE expression (Figure 2A). Interestingly, whereas SoxB1 factors inhibited expression of neural crest markers and SoxE factors promoted this expression, both inhibited expression of *epidermal keratin* (Figure 3B,C). There remains much to be learned about the distinct and overlapping regulatory activities of these two classes of Sox Factors, and how their differential deployment contributes to pluripotency and lineage restriction decisions in the early embryo.

Understanding the mechanisms underlying the evolution of gene regulatory networks is a subject of great interest and importance. Evolution of these networks has been facilitated by gene and genome duplications that increased the number and type of network components that could be deployed. The Sox superfamily has proven particularly ‘evolutionarily pliable,’ having undergone multiple rounds of duplication, divergence, sub-functionalization and neo-functionalization (Guth and Wegner, 2008; Heenan et al., 2016; Tai et al., 2016). In contrast to most neural crest potency factors (including *Snail1, Myc, Foxd3, Ets1, Ap2*, and *Vent2*)*, Sox9* and *Sox10* are not first expressed in pluripotent naïve blastula cells. SoxE expression commences in definitive neural crest cells at the neural plate border only once SoxB1 factors have become restricted to the neural progenitor pool. Our findings suggest a model in which the essential role of SoxB1 in regulating the blastula stem cell state is handed off to SoxE factors which function in part to retain the developmental potential of the neural crest. We hypothesize that this switch in Sox factor deployment facilitates the subsequent lineage restriction of neural crest cells to non-neural cell types including cartilage, melanocytes and glia.

## Material and Methods

### Embryological methods

Collection, manipulation and *in situ* hybridization of *Xenopus* embryos were performed as previously described (Bellmeyer et al. 2003) using digoxigenin-labeled RNA probes detected with BM Purple AP Substrate (Roche). mRNA for microinjection was synthesized *in vitro* from linearized plasmid templates using the SP6 Message Machine kit (Ambion). β-galactosidase mRNA was co-injected as lineage tracer and was detected with Red-Gal substrate (Research Organics). All results shown are representative of at least three biological replicates of independent experiments. Explants of naïve ectoderm were manually dissected from the animal pole of blastula (stage 8-9) embryos previously injected at two-cell stage with the indicated mRNA or morpholino. Explants were cultured at room temperature in 1X MMR on agar coated dishes until the indicated stage, and fixed in formaldehyde for 40 minutes before being processed for *in situ* hybridization. For Activin treatment of animal pole explants of naïve ectoderm were manually dissected from animal pole of blastula (stage 8-9) embryos previously injected with mRNA or morpholino. Injected explants were culture in 1X MMR. Activin prepared from R&D Systems Human/Mouse/Rat Activin A Recombinant Protein concentrated to an effective stock concentration of 20 μg/ml. Explants were treated with Activin at the stage indicated in 1X MMR supplemented with 0.1% BSA as carrier. Concentrations of 20ng/μl or 200 ng/μl were used to induce mesoderm or endoderm respectively. Explants were cultured at room temperature in 1X MMR on agar coated dishes until the indicated stage, samples were fixed in formaldehyde for 40 minutes before being processed of *in situ* hybridization.

### Dexamethasone treatment of whole embryos

Embryos were injected with Sox2-GRcMT, Sox3-GRcMT, Sox9-GRcMT, Sox10-GRcMT at two-cell stage and cultured in 0.1X MMR. Embryos were induced with a solution of 0.1X MMR with 4μg/ml Dexamethasone (Sigma) at stage 10 to bypass blastula stages. Embryos were cultured at room temperature until stage 15 (neurula) and, fixed in formaldehyde for 1 hour before being processed for *in situ* hybridization.

### Western blot analysis

For western blot, five embryos per injection condition were collected at stage 10 and lysed 1% NP-40 supplemented with a protease inhibitor cocktail (Roche), phenylmethylsulfonyl fluoride, aprotinin, and leupeptin. Proteins were detected using a primary antibody against epitope tag: Myc 1:3000 (9E10, Santa Cruz Biotechnology), and a secondary antibody conjugated to Horseradish peroxidase (HRP) and detected by chemiluminescence (GE Healthcare). Alternately, a goat anti-mouse IgG IRDye secondary was used at 1:20,000 and samples were detected via Li-COR Odyssey Imaging Systems.

## DNA constructs

The morpholino antisense oligonucleotides against the 5’UTR and coding region of *Xenopus* Sox2 (5’-GCGGAGCTACATGTCGTACTACCTC-3’), Sox3 (5’-ACTTCGAGGTTTACATATCGTACAA-3’) (Gene Tools) were previously described and validated (Schlosser et al., 2008). Embryos were injected with 5ng of each morpholino to inhibit Sox2 and Sox3 expression. The morpholino targeting Sox10 (5’-AGCTTTGGTCATCACTCATGGTGCC-3’), as described in (Taylor & LaBonne, 2005) was used to inhibit Sox10 expression (10ng). For rescue experiments, mRNA encoding epitope tagged forms of Sox2, Sox3, Sox9, or Sox10 was injected together with mRNA encoding lineage tracer β-gal. Animal caps were dissected and cultured to the indicated stage and fixed for 40 minutes in formaldehyde prior to *in situ* hybridization.

## Acknowledgements

The authors thank Joe Nguyen and Thomas Yeo for invaluable technical assistance, and members of the laboratory for helpful discussions. This work was supported by NIH R01GM116538 to CL

## Competing Interests

The authors declare no competing financial interests.

---

**Supplementary Figure 1.**
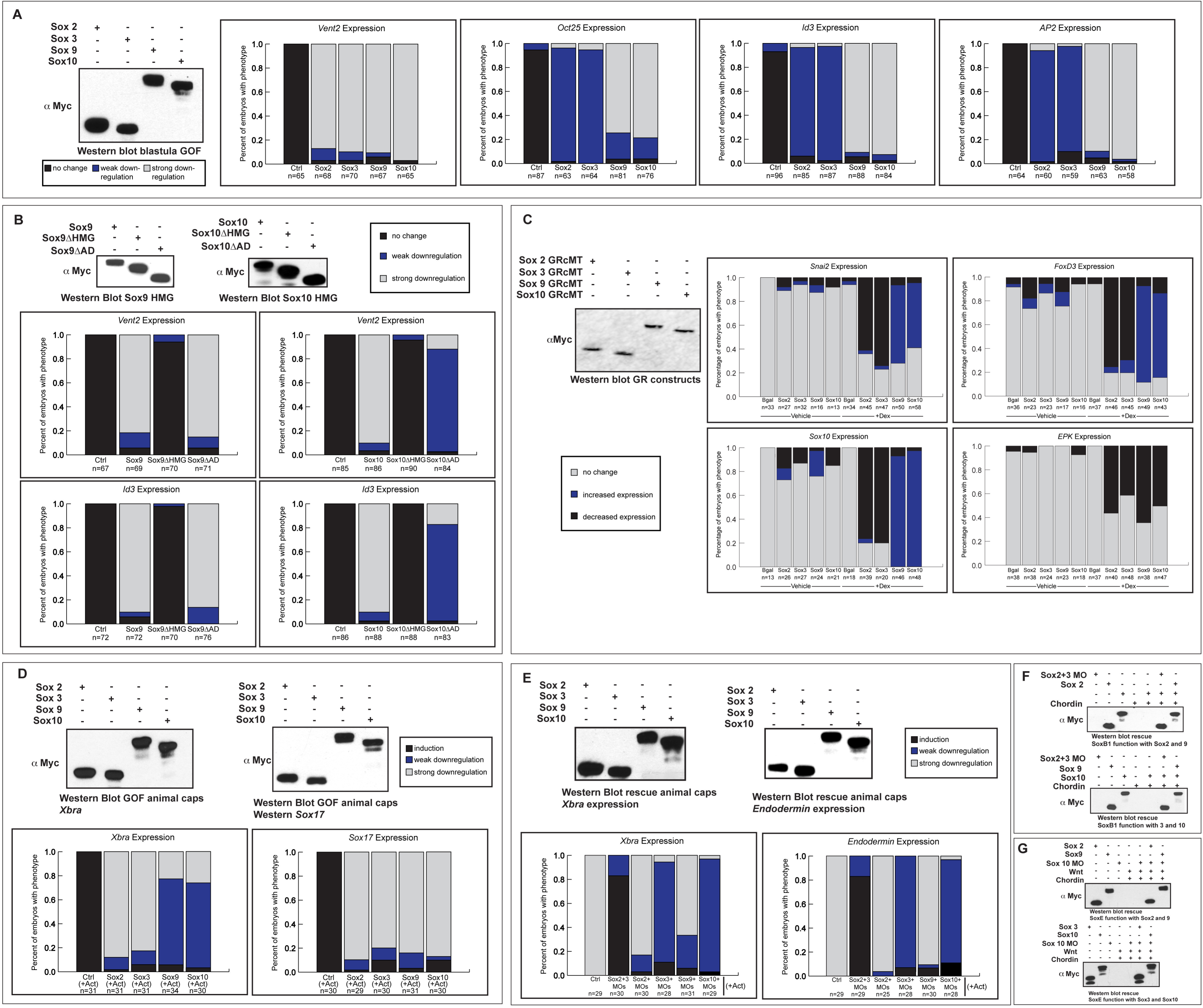
Summary of quantification for experiments and western blots to validate protein levels. (A) Western blot and *in situ* phenotype scoring associated with Figure 2A. (B) Western blot and *in situ* phenotype scoring associated with Figure 2C,D. (C) Western blot and *in situ* phenotype scoring associated with Figure 3B,C. (D) Western blot and *in situ* phenotype scoring associated with Figure 4B,C. (E) Western blot and *in situ* phenotype scoring associated with Figure 5B,C. (F) Western blot associated with Figure 6. (G) Western blot associated with Figure 7.

**Supplementary Figure 2.**
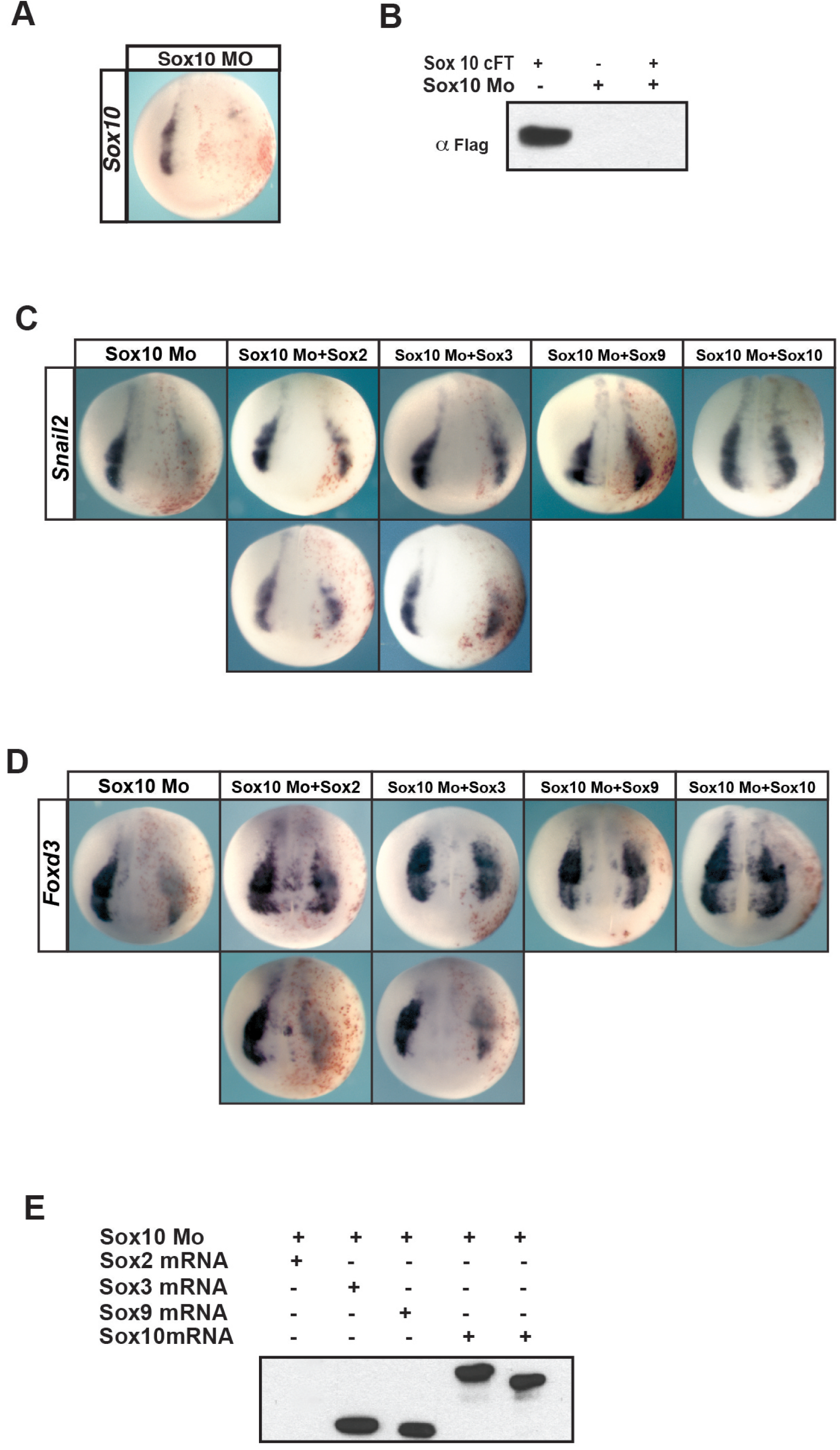
Validation of Sox10 morpholino and whole embryo rescue experiments. (A) *In situ* hybridization examining *Sox10* expression in Sox10 morpholino injected embryos. (B) Validation of Sox10 morpholino by western blot. (C) *In situ* hybridization examining *Snail2* expression in neurula (stage 15) embryos injected unilaterally with Sox10 morpholino and Sox2, Sox3, Sox9, or Sox10 mRNA. (D) (C) *In situ* hybridization examining *FoxD3* expression in neurula (stage 15) embryos injected unilaterally with Sox10 morpholino and Sox2, Sox3, Sox9, or Sox10 mRNA. (E) Western blot for Sox factor protein levels from experiments in (C, D).

## References

Abdelalim. M. E., Emara, M.M., and Kolatkar. P.R. (2014). The SOX Transcription Factors as Key Players in Pluripotent Stem Cells. Stem Cells and Development 23, 2687–2699.

Acloque H, Ocaña OH, Matheu A, Rizzoti K, Wise C, Lovell-Badge R, Nieto MA. (2011) Reciprocal repression between Sox3 and snail transcription factors defines embryonic territories at gastrulation. Dev Cell. 21(3):546–58.

Ariizumi, T., & Asashima, M. (2001). In vitro induction systems for analyses of amphibian organogenesis and body patterning., 45(1), 273–279.

Asashima, M., Nakano, H., Shimada, K., Kinoshita, K., Ishii, K., Shibai, H., & Ueno, N. (1990a). Mesodermal induction in early amphibian embryos by activin A (erythroid differentiation factor). Roux’s Archives of Developmental Biology: the Official Organ of the EDBO, 198(6), 330–335. http://doi.org/10.1007/BF00383771

Asashima, M., Nakano, H., Uchiyama, H., Davids, M., Plessow, S., Loppnow-Blinde, B., Hoppe P., Dau H., & Tiedemann H. (1990b). The vegetalizing factor belongs to a family of mesoderm-inducing proteins related to erythroid differentiation factor. Die Naturwissenschaften, 77(8), 389–391.

Akiyama, H., Chaboissier, M.-C., Martin, J. F., Schedl, A. and de Crombrugghe, B. (2002). The transcription factor Sox9 has essential roles in successive steps of the chondrocyte differentiation pathway and is required for expression of Sox5 and Sox6. Genes Dev. 16, 2813–2828.

Aoki, Y., Saint-Germain, N., Gyda, M., Magner-Fink, E., Lee, Y.-H., Credidio, C. and Saint-Jeannet, J.-P. (2003). Sox10 regulates the development of neural crest-derived melanocytes in Xenopus. Developmental Biology 259, 19–33.

Avilion AA, Nicolis SK, Pevny LH, Perez L, Vivian N, Lovell-Badge R. (2003) Multipotent cell lineages in early mouse development depend on SOX2 function. Genes Dev. 17(1):126–40.

Bellmeyer, A., Krase, J., Lindgren, J. and LaBonne, C. (2003). The Protooncogene c-Myc Is an Essential Regulator of Neural Crest Formation in Xenopus. Developmental Cell 4, 827–839.

Bergsland M, Werme M, Malewicz M, Perlmann T, Muhr J. (2006) The establishment of neuronal properties is controlled by Sox4 and Sox11. Genes Dev. 20(24):3475–86.

Bergsland, M., Ramsköld, D., Zaouter, C., Klum, S., Sandberg, R. and Muhr, J. (2011). Sequentially acting Sox transcription factors in neural lineage development. Genes Dev. 25, 2453–2464.

Bi, W., Deng, J. M., Zhang, Z., Behringer, R. R. and de Crombrugghe, B. (1999). Sox9 is required for cartilage formation. Nat. Genet. 22, 85–89.

Boer, J. Kopp, S. Mallanna, M. Desler, H. Chakravarthy, P.J. Wilder, C. Bernadt, A. Rizzino (2007) Elevating the levels of Sox2 in embryonal carcinoma cells and embryonic stem cells inhibits the expression of Sox2:Oct-3/4 target genes. Nucleic Acids Res., 35. 1773–1786

Bowles, J., Schepers, G. and Koopman, P. (2000). Phylogeny of the SOX family of developmental transcription factors based on sequence and structural indicators. Developmental Biology 227, 239–255.

Britsch S, Goerich DE, Riethmacher D, Peirano RI, Rossner M, Nave KA, Birchmeier C, Wegner M. (2001) The transcription factor Sox10 is a key regulator of peripheral glial development. Genes Dev. 15(1):66–78.

Buitrago-Delgado, E., Nordin, K., Rao, A., Geary, L. and LaBonne, C. (2015). Shared regulatory programs suggest retention of blastula-stage potential in neural crest cells. Science 348, 1332–1335.

Charney RM, Forouzmand E, Cho JS, Cheung J, Paraiso KD, Yasuoka Y, Takahashi S, Taira M, Blitz IL, Xie X, Cho KW. (2017) Foxh1 Occupies cis-Regulatory Modules Prior to Dynamic Transcription Factor Interactions Controlling the Mesendoderm Gene Program. Dev Cell. 40(6):595–607.

Chen, X., Xu, H., Yuan, P., Fang, F., Huss, M., Vega, V. B., Wong, E., Orlov, Y. L., A., Birchmeier, C. and Wegner, M. (2001). The transcription factor Sox10 is a key regulator of peripheral glial development. Genes Dev. 15, 66–78.

Cheung, M. and Briscoe, J. (2003). Neural crest development is regulated by the transcription factor Sox9. Development 130, 5681–5693.

Geary, L and LaBonne, C. (2018) FGF mediated MAPK and PI3K/Akt signals make distinct contributions to pluripotency and the establishment of neural crest. Elife 7, 139–159

Guth, S. I. E. and Wegner, M. (2008). Having it both ways: Sox protein function between conservation and innovation. Cell. Mol. Life Sci. 65, 3000–3018.

Graham, V., Khudyakov, P., Ellis, P., Pevny, L. (2003). Sox2 functions to maintain neural progenitor identity. Neuron. 39.5. 749–765.

Haldin, C. E. and LaBonne, C. (2010). SoxE factors as multifunctional neural crest regulatory factors. The International Journal of Biochemistry & Cell Biology 42, 441–444.

Hall, B. K. (2009). Embryological Origins and the Identification of Neural Crest Cells. In The Neural Crest and Neural Crest Cells in Vertebrate Development and Evolution, pp. 23–61. Springer, Boston, MA.

Hall, B. K. (2013). The Neural Crest in Development and Evolution. New York, NY: Springer Science & Business Media.

Heenan, P., Zondag, L. and Wilson, M. J. (2016). Evolution of the Sox gene family within the chordate phylum. Gene 575, 385–392.

His, W. Untersuchungen über die erste Anlage des Wirbeltierleibes: die erste Entwickelung des Hühnchens im Ei, Vogel FCW, Leipzig. (1868).

Hoffmann SA, Hos D, Küspert M, Lang RA, Lovell-Badge R, Wegner M, Reiprich S. (2014) Stem cell factor Sox2 and its close relative Sox3 have differentiation functions in oligodendrocytes. Development. 141(1):39–50.

Hong, C.-S. and Saint-Jeannet, J.-P. (2005). Sox proteins and neural crest development. Semin. Cell Dev. Biol. 16, 694–703.

Hong, C.-S. and Saint-Jeannet, J.-P. (2007). The activity of Pax3 and Zic1 regulates three distinct cell fates at the neural plate border. Molecular Biology of the Cell 18, 2192–2202.

Honoré SM1, Aybar MJ, Mayor R. (2003) Sox10 is required for the early development of the prospective neural crest in Xenopus embryos. Dev Biol. 260(1):79–96.

Hoser M, Potzner MR, Koch JM, Bösl MR, Wegner M, Sock E. (2008) Sox12 deletion in the mouse reveals nonreciprocal redundancy with the related Sox4 and Sox11 transcription factors. Mol Cell Biol. 28(15):4675–87.

Kelsh, R. N. (2006). Sorting out Sox10 functions in neural crest development. Bioessays 28, 788–798.

Kim, J., Lo, L., Dormand, E. and Anderson, D. J. (2003). SOX10 maintains multipotency and inhibits neuronal differentiation of neural crest stem cells. Neuron 38, 17–31.

Kopp, JJ, Ormsbee, BD, Desler, M. Rizzino, A (2008) Small increases in the level of Sox2 trigger the differentiation of mouse embryonic stem cells Stem Cells, 26 903–911.

LaBonne, C., & Bronner-Fraser, M. (1998). Neural crest induction in Xenopus: evidence for a two-signal model. Development (Cambridge, England), 125(13), 2403–2414.

Le Douarin, N. and Kalcheim, C. (2009). The Neural Crest. 2nd ed. Cambridge: Cambridge University Press.

Lee, P.-C., Taylor-Jaffe, K. and LaBonne, C. (2011). SUMO regulation of SoxE factors during neural crest development. Developmental Biology 356, 259.

Lee, P.-C., Taylor-Jaffe, K.M., Nordin, K.M., Prasad, M.S., Lander, R. M. and LaBonne, C. (2012). SUMOylated SoxE factors recruit Grg4 and function as transcriptional repressors in the neural crest. J. Cell Biol. 198, 799–813.

Lefebvre V, Li P, de Crombrugghe B. (1998) A new long form of Sox5 (L-Sox5), Sox6 and Sox9 are coexpressed in chondrogenesis and cooperatively activate the type II collagen gene. EMBO J. 117(19):5718–33.

Lefebvre, V. and Dvir-Ginzberg, M. (2017). SOX9 and the many facets of its regulation in the chondrocyte lineage. Connect. Tissue Res. 58, 2–14.

Light, W., Vernon, A, Lasorella, A., Iavarone, A. and LaBonne, C. (2005). Xenopus Id3 is required downstream of Myc for the formation of multipotent neural crest progenitor cells. Development 132, 1831–1841.

Masui, S., Nakatake, Y., Toyooka, Y., Shimosato, D., Yagi, R., Takahashi, K., Okochi, H., Okuda, A., Matoba, R., Sharov, A. A., et al. (2007). Pluripotency governed by Sox2 via regulation of Oct3/4 expression in mouse embryonic stem cells. Nat. Cell Biol. 9, 625–635.

Mori-Akiyama, Y., Akiyama, H., Rowitch, DH. and de Crombrugghe, B. (2003). Sox9 is required for determination of the chondrogenic cell lineage in the cranial neural crest. Proc. Natl. Acad. Sci. U.S.A. 100, 9360–9365.

Nordin, K. and LaBonne, C. (2014). Sox5 Is a DNA-binding cofactor for BMP R-Smads that directs target specificity during patterning of the early ectoderm. Developmental Cell 31, 374–382.

Ochoa, S. Salvador, S, LaBonne, C (2012). The LIM adaptor protein LMO4 is an essential regulator of neural crest development. Developmental Biology 361:313–325.

O’Donnell, M., Hong, C.-S., Huang, X., Delnicki, R. J. and Saint-Jeannet, J.-P. (2006). Functional analysis of Sox8 during neural crest development in Xenopus. Development 133, 3817–3826.

Prasad, M. S., Sauka-Spengler, T. and LaBonne, C. (2012). Induction of the neural crest state: control of stem cell attributes by gene regulatory, post-transcriptional and epigenetic interactions. Developmental Biology 366, 10–21.

Rao and C. LaBonne. (2018) Regulation of Histone Acetylation controls the transition from plutipotency to lineage restriction in early embryos. (Development, in press)

Rex M, Orme A, Uwanogho D, Tointon K, Wigmore PM, Sharpe PT, Scotting PJ. (1997) Dynamic expression of chicken Sox2 and Sox3 genes in ectoderm induced to form neural tissue. Dev Dyn. 209(3):323–32.

Saint-Germain N, Lee YH, Zhang Y, Sargent TD, Saint-Jeannet JP. (2004) Specification of the otic placode depends on Sox9 function in Xenopus. Development. 131(8):1755–63.

Sarkar, A. and Hochedlinger, K. (2013). The sox family of transcription factors: versatile regulators of stem and progenitor cell fate. Cell Stem Cell 12, 15–30.

Sasai, Y., Lu, B., Steinbeisser, H., & De Robertis, E. M. (1995). Regulation of neural induction by the Chd and Bmp-4 antagonistic patterning signals in Xenopus., 376(6538), 333–336. http://doi.org/10.1038/376333a0

Schepers, G. E., Teasdale, R. D. and Koopman, P. (2002). Twenty pairs of sox: extent, homology, and nomenclature of the mouse and human sox transcription factor gene families. Developmental Cell 3, 167–170.

Schlosser G, Awtry T, Brugmann SA, Jensen ED, Neilson K, Ruan G, Stammler A, Voelker D, Yan B, Zhang C, Klymkowsky MW, Moody SA. (2008) Eya1 and Six1 promote neurogenesis in the cranial placodes in a SoxB1-dependent fashion. Dev Biol.; 320(1):199–214.

Spokony, R. F., Aoki, Y., Saint-Germain, N., Magner-Fink, E. and Saint-Jeannet, J.-P. (2002). The transcription factor Sox9 is required for cranial neural crest development in Xenopus. Development 129, 421–432.

Stolt CC, Schlierf A, Lommes P, Hillgärtner S, Werner T, Kosian T, Sock E, Kessaris N, Richardson WD, Lefebvre V, Wegner M. (2006) SoxD proteins influence multiple stages of oligodendrocyte development and modulate SoxE protein function. Dev Cell. 11(5):697–709.

Stolt CC, Lommes P, Hillgärtner S, Wegner M. (2008) The transcription factor Sox5 modulates Sox10 function during melanocyte development. Nucleic Acids Res. 36(17):5427–40.

Streit A, Sockanathan S, Pérez L, Rex M, Scotting PJ, Sharpe PT, Lovell-Badge R, Stern CD. (1997) Preventing the loss of competence for neural induction: HGF/SF, L5 and Sox-2. Development. 124(6):1191–202.

Suzuki, T., Sakai, D., Osumi, N., Wada, H. and Wakamatsu, Y. (2006). Sox genes regulate type 2 collagen expression in avian neural crest cells. Dev. Growth Differ. 48, 477–486.

Tai, A., Cheung, M., Huang, Y.-H., Jauch, R., Bronner, M. E. and Cheah, K. S. E. (2016). SOXE neofunctionalization and elaboration of the neural crest during chordate evolution. Sci Rep 6, 34964.

Takahashi, K. and Yamanaka, S. (2006). Induction of pluripotent stem cells from mouse embryonic and adult fibroblast cultures by defined factors. Cell 126, 663–676.

Taylor, K. M. and LaBonne, C. (2005). SoxE Factors Function Equivalently during Neural Crest and Inner Ear Development and Their Activity Is Regulated by SUMOylation. Developmental Cell 9, 593–603.

Taylor, K. M. and LaBonne, C. (2007). Modulating the activity of neural crest regulatory factors. Curr. Opin. Genet. Dev. 17, 326–331.

Thomson, M., Liu, S. J., Zou, L.-N., Smith, Z., Meissner, A. and Ramanathan, S. (2011). Pluripotency factors in embryonic stem cells regulate differentiation into germ layers. Cell 145, 875–889.

Wegner, M. (1999). From head to toes: the multiple facets of Sox proteins. Nucleic Acids Res. 27, 1409–1420.

Wegner, M. (2011). SOX after SOX: SOXession regulates neurogenesis. Genes Dev. 25, 2423–2428.

Xu J, Watts JA, Pope SD, Gadue P, Kamps M, Plath K, Zaret KS, Smale ST. (2009). Transcriptional competence and the active marking of tissue-specific enhancers by defined transcription factors in embryonic and induced pluripotent stem cells. Genes Dev. 23(24):2824–38.

Yamaguchi S, Hirano K, Nagata S, Tada T. (2011) Sox2 expression effects on direct reprogramming efficiency as determined by alternative somatic cell fate. Stem Cell Res. 2011 Mar;6(2): 177–86.

